# Antiviral effect of a nucleic acid hydrolyzing scFv against oseltamivir resistant influenza A virus

**DOI:** 10.1101/2021.02.19.432068

**Authors:** Yongjun Lee, Dongjun Kim, Taehyun Kim, Yeonsu Oh, Won-Keun Kim, Sukchan Lee

## Abstract

Influenza viral genome is frequently mutated due to antigenic shift and drift, and therefore the existing antiviral drugs have been suffered from low efficacy against the viruses. Here we report an innovative strategy for treating influenza Type A (IAV) infection by 3D8 single chain variable fragment (scFv) showing intrinsic viral RNA hydrolyzing activity, cell penetration activity and the epidermal cell penetration ability. In this study, we first analyzed antiviral activity directed by 3D8 scFv to three different strains, two oseltamivir-sensitive (A/PuertoRico/8/1934, A/X-31) and oseltamivir-resistant (A/Korea/2785/2009pdm) using cell culture models, suggesting that 3D8 scFv reduces viral genomic RNA. Moreover, we further addressed antiviral effect to analyze clinical outcomes in *in vivo* mice model. Intranasal administered 3D8 scFv rescued the mice challenged by oseltamivir resistant H1N1. Consistent results are observed through IHC pathology and molecular virological analysis. Taken together, these results demonstrate that 3D8 scFv has antiviral potential against a wide range of IAV.

## Introduction

Influenza, commonly known as ‘the flu’, is an infectious pyrogenic respiratory disease caused by an influenza virus. Annual outbreaks are reported globally and have significant health, social, and economic impacts on affected communities (1–3). There are two main types of influenza virus: Types A and B, which routinely spread in people are responsible for seasonal flu epidemics each year. Influenza spreads around the world in yearly outbreaks, resulting in about three to five million cases of severe illness and about 290,000 to 650,000 deaths according to the World Health Organization (WHO). Yearly vaccinations against influenza are recommended by the WHO for those at high risk and the vaccine is usually effective against three or four types of influenza. However, a vaccine made for one year may not be useful in the following year, since the virus evolves rapidly. Influenza A and B viruses can change in two different ways-antigenic drift and antigenic shift (4, 5).

Influenza A (H1N1/pdm09) virus (IAV) that emerged in Mexico and United States, April 2009 caused the first global influenza pandemic in more than 40 years. This new virus contained a unique combination of influenza genes not previously identified in animals or people. Disease burden of the IAV H1N1/pdm09 continued for the next 10 years. The Center for Disease Control and Prevention (CDC) estimated that 10.5 million people were infected and 75,000 people died from the IAV H1N1/pdm09 infection (6–8).

Antiviral medication such as neuraminidase inhibitor oseltamivir (Osel), among others, have been used to treat influenza. Oseltamivir is the most widely used antiviral drug targeting NA proteins and consequently suppresses viral proliferation (9, 10).

However, the IAV H1N1/pdm09 is found to be resistant to oseltamivir through a particular genetic change known as the ‘H275Y’ mutation. The ‘H275Y’ mutation makes oseltamivir ineffective in treating illnesses with the virus (11, 12). In addition, antiviral drugs of other mechanisms against IAV, such as M2 ion channel inhibitor like amantadine and rimantadine, may also be less effective at any time due to such frequent viral changes.

3D8 scFv is a recombinant single chain antibody (~28 kDa) developed by linking the variable heavy (VH) and light (VL) chain domains with a flexible peptide linker. The origin of 3D8 scFv was found in MRL-*lpr/lpr* mice, which developed an autoimmune syndrome that was similar to human systemic lupus erythematosus (13, 14). 3D8 scFv has been known to penetrate various cell types through caveolae-mediated endocytosis without any carrier (15). Moreover, 3D8 scFv has intrinsic nucleic acid hydrolyzing activity and can directly breakdown viral RNA in a sequence independent manner but exhibited rare cytotoxicity at specific protein concentration (16). Previous studies have demonstrated the antiviral effect of 3D8 scFv against a broad spectrum of viruses including the classical swine fever virus, human herpes simplex virus and pseudorabies virus (17, 18). While oseltamivir is unable to block NA in viruses with specific mutations in this protein indicating, and is ineffective as an antiviral drug, 3D8 scFv breaks down the viral RNA in the cytoplasm regardless of the virus type; thus, it is predicted to exhibit antiviral effect even in mutated viruses.

In this study, it was found that 3D8 scFv exhibits antiviral effect on oseltamivir-sensitive influenza A/PuertoRico/8/1934 (H1N1/PR8) and A/X-31 (H3N2/X-31) and oseltamivir-resistant influenza A/Korean/2785/2009pdm (H1N1/H275Y) viruses. The results validate the effectiveness of this new treatment that differs from the current standard drugs used for the prevention and therapeutics against IAV infection including oseltamivir-resistant strains.

## Results

### 3D8 scFv increases the viability of cells infected with three different influenza A viruses

An important prerequisite for the antiviral effect of 3D8 scFv is probably the ability to penetrate cells of various types and origins. Human (A549 and HeLa), simian (Vero), and canine (Madin–Darby canine kidney; MDCK) cell lines were used to prove this. The 3D8 scFv was detected in the cytoplasm of all cell lines within 24h post treatment (Fig. 1*A*). To examine the ability of 3D8 scFv to hydrolize viral RNA, the cells treated with 3D8 scFv (0.5 μg) were mixed with viral NA or nucleoprotein (NP) RNA (1 μg each) from H1N1/PR8 and incubated at 37°C, and then viral RNA was analyzed through agarose gel running. The results showed that the viral RNA was disintegrated (Fig. 1*B*). Next, MDCK cells were treated with different concentrations of 3D8 scFv (0 to 10 μM) to confirm its cytotoxicity. A minimum cytotoxicity of less than 5% was observed across all treatments, suggesting that 3D8 scFv dependent cytotoxicity remained minimal at tested working concentration (Fig. 1*C*).

**Fig. 1.**
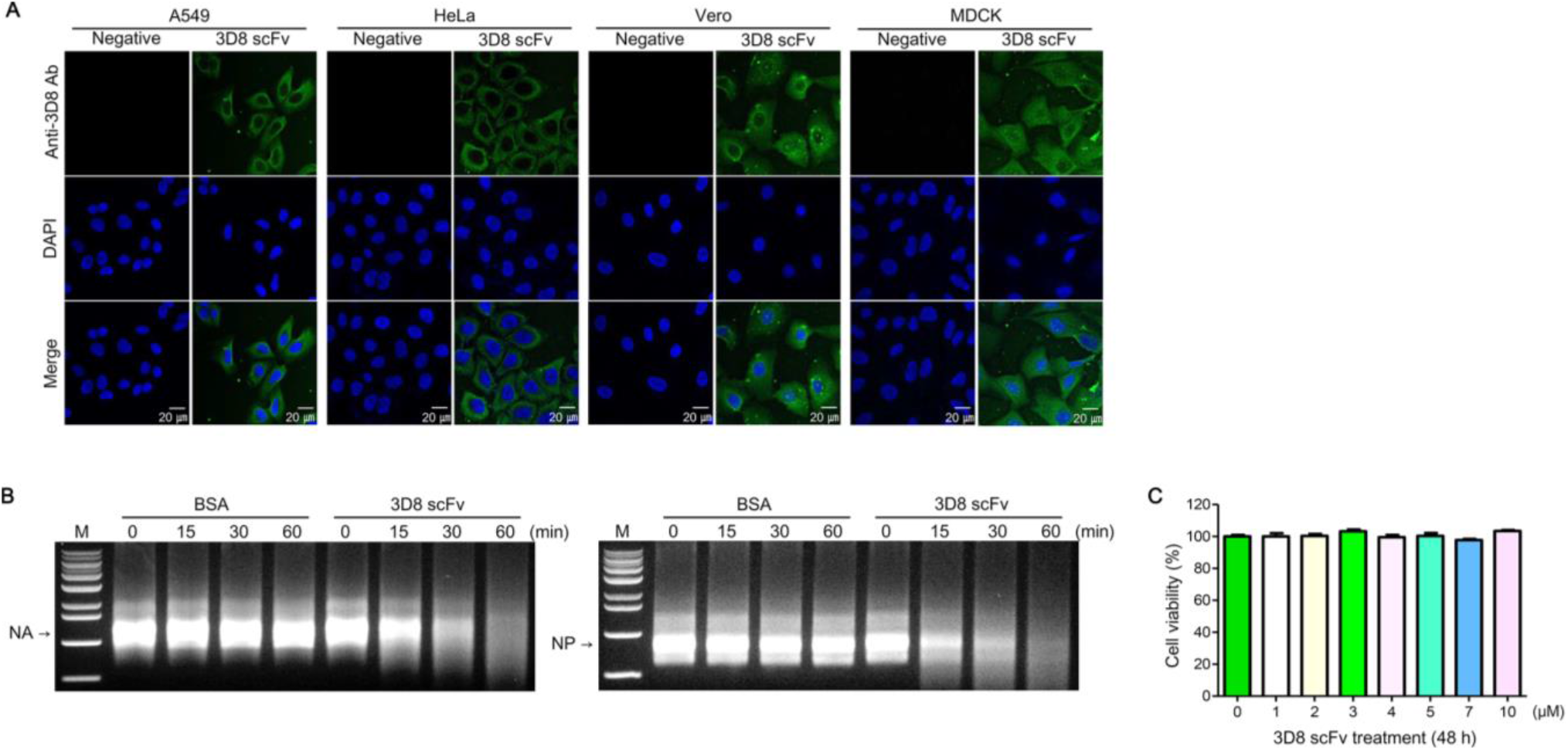
Characteristics of 3D8 scFv. (*A*) Cell penetration ability of 3D8 scFv was assessed using immunocytochemistry in A549, HeLa, Vero, and MDCK cells. Cells were incubated with 3D8 scFv for 24 h at 37℃. Nuclei were detected by DAPI staining (blue), 3D8 scFv was visualized using antibody conjugated with Alexa flour 488 fluorescent dye (Green). Scale bars represent 20 μm. (*B*) Viral RNA (NA and NP) hydrolyzing activity of 3D8 scFv was examined by agarose gel electrophoresis. 3D8 scFv was incubated with the synthesized viral RNA in TBS buffer for 1 h at 37℃. (*C*) Cytotoxicity of 3D8 scFv in MDCK cells. MDCK cells were incubated in the presence of various concentrations of 3D8 scFv (0 to 10 μM) for 48 h at 37℃. Cell viability was measured using the MTT assay.

After the cell line was infected with three different viral subtypes, H1N1/PR8, H3N2/X-31, and H1N1/H275Y, the viability between cells treated with and without 3D8 scFv was compared using MTT assay. When cells infected with each of the three types of viruses (MOI = 0.1) were treated with different concentrations of oseltamivir, the H1N1/H275Y-infected cells exhibited drug resistant phenotype but not the H1N1/PR8- and H3N2/X-31-infected cells (*SI Appendix*, Fig. S1). To determine the antiviral effect of 3D8 scFv treated at pre- or post-viral infection, MDCK cells were treated with three different concentrations (3, 5, and 7 μM) of 3D8 scFv 24 h prior to viral infection and incubated for additional 48 h (Fig. 2*A*). Alternatively, the cells were treated with 3D8 scFv at 1 h post viral infection and incubated for 47 h (Fig. 2*E*). In the 3D8 scFv pre-treatment group, the viability of H1N1/PR8 (0 μM) infected cells was initially 66.8%, and it increased by 20% to 88.3% after treatment with 5 μM 3D8 scFv (Fig. 2*B*). Moreover, while the viability of H3N2/X-31 (0 μM) infected cells was 54.5%, treatment with 3 μM of 3D8 scFv increased the viability by 20% to 74.4% (Fig. 2*C*). Furthermore, the viability of all 3D8 scFv treatment groups in the H1N1/H275Y-infected cells increased by at least 20% to 84.8% compared to the non 3D8 scFv-treated level (55.6% [0 μM]; Fig. 2*D*). In 3D8 scFv post-treatment, while H1N1/PR8 (0 μM) infected cell viability was 63.2%, treatment with 5 μM 3D8 scFv increased the viability by 23% to 86.9% (Fig. 2*F*). Moreover, while the viability of H3N2/X-31 (0 μM) infected cells was 62.7%, treatment with 5 μM of 3D8 scFv increased the viability by 22% to 84.4% (Fig. 2*G*). Further, the treatment of H1N1/H275Y-infected cells with 3D8 scFv increased the viability by 20% to 77.4% compared with that of the non 3D8 scFv-treated cells (58.4% [0 μM]; Fig. 2*H*). It was resulted that pre-/post-treatments with 3D8 scFv *in vitro* increased the viability of virus-infected cells by 20%.

**Figure 2.**
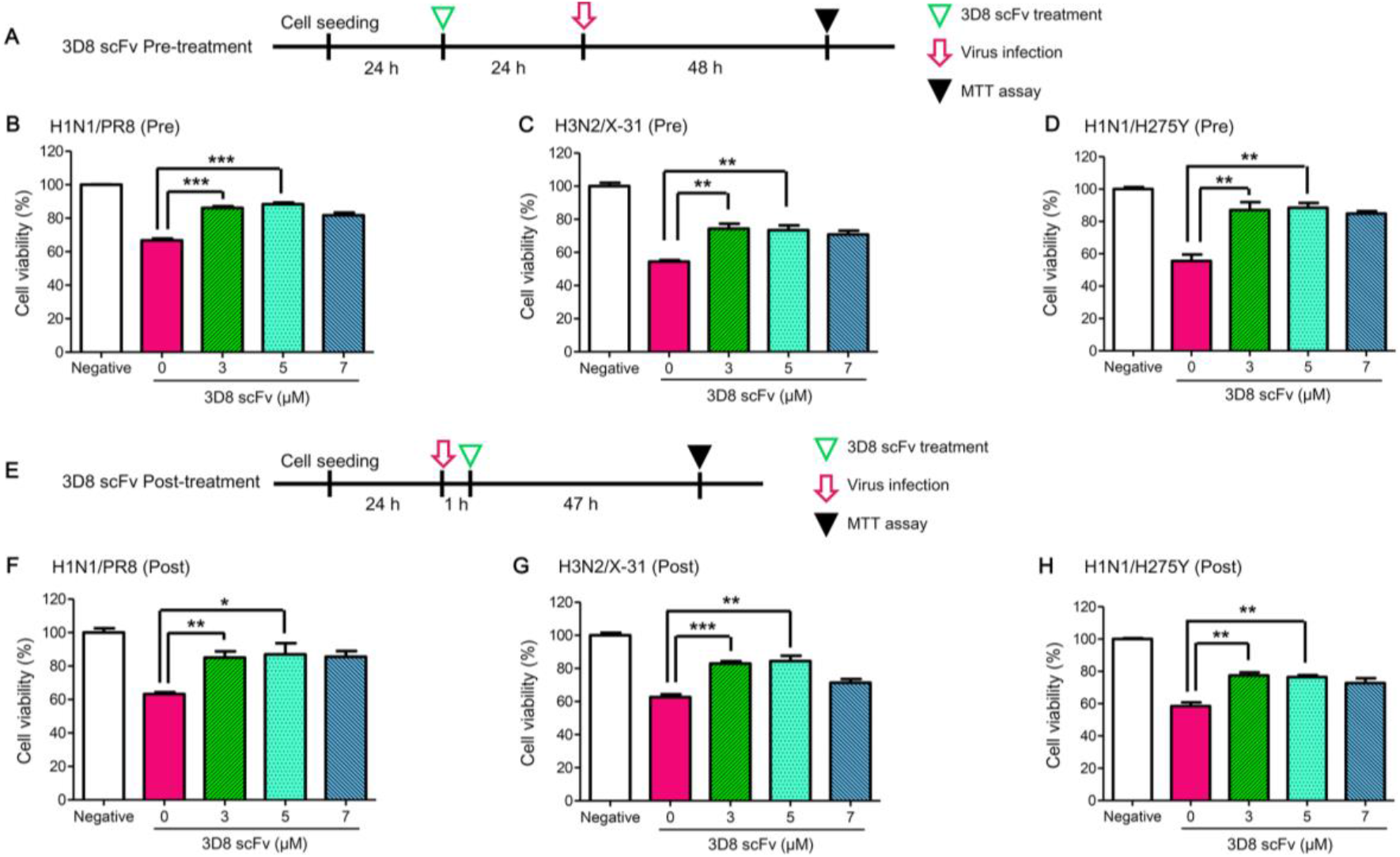
Dose dependent antiviral activity of 3D8 scFv against three influenza virus strains. (*A* and *E*) Schematic diagram for testing the antiviral effect (cell viability) of 3D8 scFv pre-/post-treatment against three viral strains (H1N1/PR8, H3N2/X-31, and H1N1/H275Y). (*B*-*D*) Relative viability of 3D8 scFv pre-treated MDCK cells against viruses. MDCK cells were incubated with 3D8 scFv (3, 5, 7 μM) at 37℃ for 24 h, and then infected with viruses (MOI = 0.1) for 1 h. The viral inoculum was replaced with new media after 1 h. After incubation at 37℃ for 48 h, cell viability was measured using the MTT assay. (*F*-*H*) Relative viability of 3D8 scFv post-treated MDCK cells against viruses. MDCK cells were infected with viruses (MOI = 0.1) for 1 h and the viral inoculum was replaced with new media containing 3D8 scFv (3, 5, 7 μM) after 1 h. Then, after incubation at 37℃ for 47 h, cell viability was measured using MTT assay. Data are shown as mean ± S.E.M of triplicate samples. Error bars indicate standard error (SE). Asterisks indicate significant difference determined by the *t*-test (*P < 0.05, **P < 0.01, ***P < 0.001).

### 3D8 scFv decreases viral RNA/protein expression

A previous study demonstrated that 3D8 scFv was able to penetrate cells within 24 h (*15*). This time, it was examined that the 3D8 scFv in virus-infected cells could inhibit viral RNA expression. MDCK cells were infected with three different viruses, and pre- and post-treatment with 3 μM of 3D8 scFv was performed, followed by measuring the expression of the viral RNAs (HA and NP) at 24 h post viral challenge (Fig. 3*A*). In H1N1/PR8-infected cells, the viral RNA expression decreased to 47% (HA) and 49% (NP) in the pre-treatment group (Fig. 3*B*), and 60% (HA) and 52% (NP) in the post-treatment cells (Fig. 3*E*) compared with the non-treated control. In H3N2/X-31-infected cells, the viral RNA expression decreased to 59% (HA) and 51% (NP) in the pre-treatment group (Fig. 3*C*) and to 55% (HA) and 58% (NP) in the post-treatment group (Fig. 3*F*). Furthermore, in the H1N1/H275Y-infected cells, the viral RNA expression decreased to 63% (HA) and 51% (NP) in the pre-treatment group and to 56% (HA) and 48% (NP) in the post-treatment group (Fig. 3 *D* and *G*).

**Figure 3.**
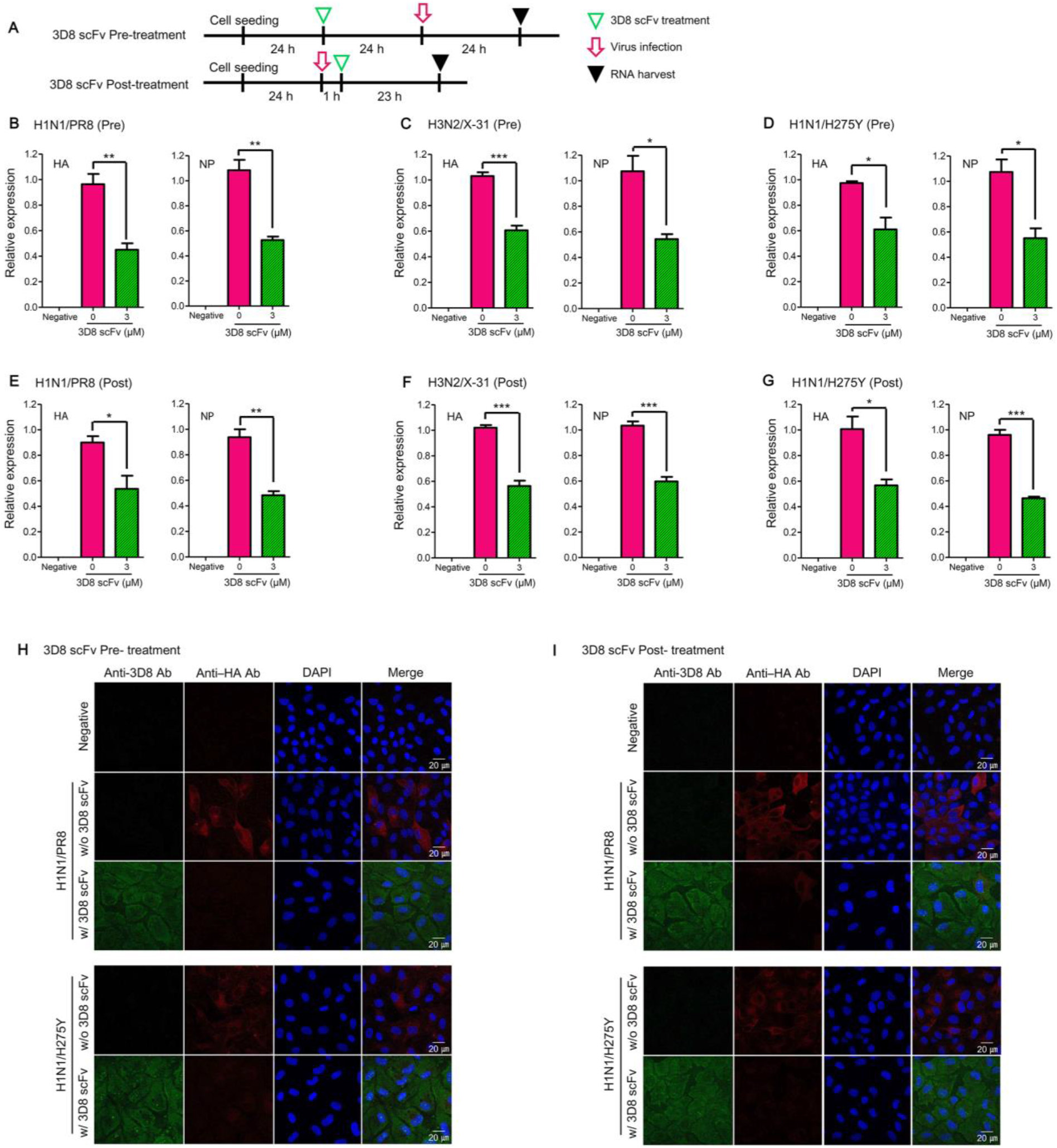
Inhibition of virus proliferation by 3D8 scFv. (*A*) Schematic diagram for antiviral test (viral RNA/protein) of 3D8 scFv pre-/post-treatment against viruses (H1N1/PR8, H3N2/X-31, and H1N1/H275Y). (*B*-*D*) Relative viral RNA expression (HA, NP) in 3D8 scFv pre-treated MDCK cells. MDCK cells were incubated with 3D8 scFv (3 μM), infected with viruses (MOI = 0.1), and qRT-PCR was performed to measure the relative expression of viral RNAs as described in materials and methods. (*E*-*G*) Relative viral RNA expression in 3D8 scFv post-treated MDCK cells. MDCK cells were infected with viruses (MOI = 0.1), treated with 3D8 scFv (3 μM), and qRT-PCR was performed to measure the relative expression of viral RNAs as described in the materials and methods. Data are shown as mean ± S.E.M of triplicate samples. Error bars indicate SE. *P < 0.05, **P < 0.01, ***P < 0.001. (*H* and *I*) Detection of viral protein (HA) in infected MDCK cells using immunocytochemistry. 3D8 scFv treatment and viral infection was the same as described in the previous section. Nuclei were detected using DAPI (Blue), 3D8 scFv with Alexa flour 488 (Green), and viral HA with TRITC (Red). Scale bars represent 20 μm.

To confirm the reduction at the viral protein level, MDCK cells were infected with H1N1/PR8 or H1N1/H275Y, and pre-/post-treated with 3D8 scFv. The cells were incubated for 24 h, and the presence of viral HA protein and 3D8 scFv was verified using immunofluorescence assay. In the majority of H1N1/PR8-infected cells without 3D8 scFv treatment, viral HA protein was detected in the cytoplasm (red signal); however, in the cells with 3D8 scFv pre-treatment, the viral protein expression was diminished compared with the non-treated control (Fig. 3*H*, upper panel). Of note, 3D8 scFv was found in the cytoplasm (green signal) in all 3D8 scFv treatment groups. In the H1N1/H275Y-infected and 3D8 scFv-treated cells, the viral HA protein signal appeared to be less than the virus infected and non-treated cells (Fig. 3*H*, lower panel). Being consistent with 3D8 scFv pre-treatment groups, the viral HA was decreased in all virus infected and 3D8 scFv post-treated groups compared with the infection-only group (Fig. 3*I*).

### 3D8 scFv increases the survival of H1N1/H275Y-infected mice

To evaluate clinical antiviral effects of 3D8 scFv, mice as an experimental *in vivo* model were tested. For the safety, mice did not show any abnormal clinical sign after 3D8 scFv was administered for 14 days (*SI Appendix,* Fig. S2*A*). Histopathologically, there is no abnormal change in the lung parenchyma and its interstitium (*SI Appendix,* Fig. S2 *B* and *C*). Of note, 3D8 scFv was found to be localized in the epithelial lining of bronchiole and alveoli for 48 h after administration (*SI Appendix,* Fig. S2 *D* and *E*).

For the efficacy, five groups of six-week-old male Balb/c mice were challenged with 8 to 10 mice per group intranasally with H1N1/H275Y, 50% lethal dose (LD_50_, 5 × 10^4^ PFU), and PBS (pre/post)-, 3D8 scFv (pre/post)- and oseltamivir-treatments according to the manufacturer’s instruction were performed, respectively. Clinical signs were monitored for the next 11 days including body weight change, coughing, hunched back, coarse fur, gathering in clusters (fever), abnormal discharge, survival, and so on (Fig. 4 *A* and *B*). The result of the survival rate was revealed that 25% of the PBS pre-treated group survived, while 90% of the 3D8 scFv pre-treatment group survived (Fig. 4*C*). 40% of the 3D8 scFv post-treatment group survived, and none of the PBS post-treated group survived (Fig. 4*E*). 30% of the oseltamivir treatment group survived.

**Figure 4.**
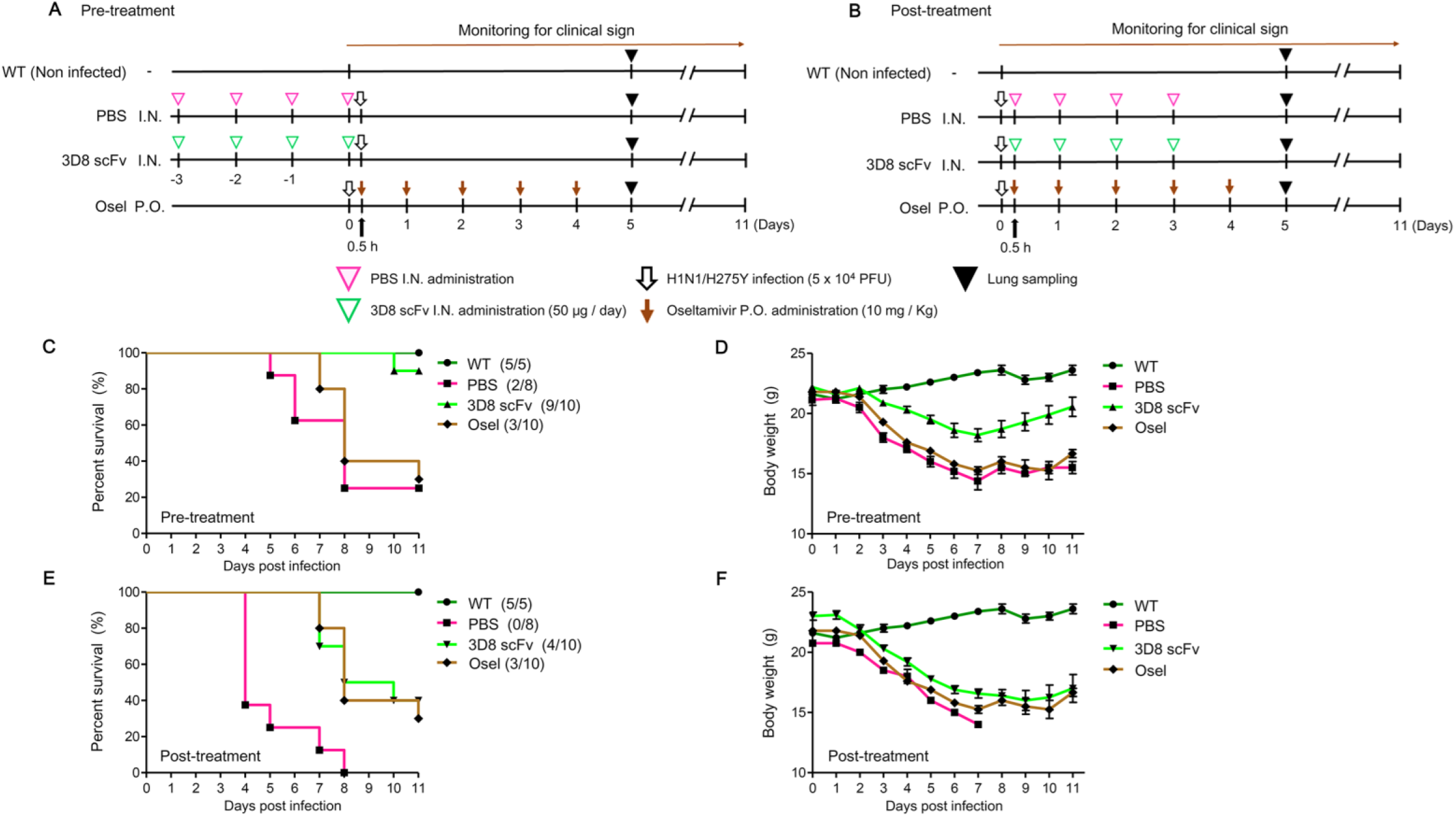
*In vivo* assessment of antiviral effects of 3D8 scFv against H1N1/H275Y infection. (*A* and *B*) Schematic diagram of 3D8 scFv treatment and H1N1/H275Y infection protocol. Mice were infected with H1N1/H275Y virus and pre-/post-treated intranasally with 3D8 scFv (50 μg/day) for 4 days. Oseltamivir (Osel) (10 mg/Kg) was orally administered (5 days consecutively). The clinical signs in the mice were observed for 11 days. The mice were euthanized (5 dpi) for lung sampling. (*C* and *E*) Survival rate of infected mice. (*D* and *F*) The mean body weight of infected mice. (WT n = 5, PBS Pre-/Post-treatment n = 8, 3D8 scFv Pre-/Post-treatment n = 10, Oseltamivir administration n = 10).

For the body weight change, in the PBS pre-treatment group, the average body weight was 21.1 g at 0 day post infection (dpi), and decreased by 27% to 15.5 g at 11 dpi (Fig. 4*D*). In the 3D8 scFv pre-treatment group, the average body weight was 22.2 g at 0 dpi, and decreased by about 18% to 18.2 g at 7 dpi. However, the average body weight rebounded to 20.6 g at 11 dpi. In the oseltamivir group, the average body weight was 21.8 g at 0 dpi, decreased by 30% at 7 dpi, and rebounded at 11 dpi.

In the PBS post-treatment group, the average body weight was 20.8 g at 0 dpi, but none of the group survived at 8 dpi (Fig. 4*E*). In the 3D8 scFv post-treatment group, the average body weight was 23 g at 0 dpi, decreased by about 30% to 16 g at 10 dpi (Fig. 4*F*). However, the average body weight restored later. These results demonstrate that 3D8 scFv had a durable clinical benefit, which consequently enhanced the survival rate and although it could not prevent initial weight loss due to viral challenge, the body weight was restored quickly.

### 3D8 scFv administration lowered histopathological lesions in the lungs of viral challenged mice

Lung tissue samples were collected at 5 dpi of H1N1/H275Y challenge (Fig. 4 *A* and *B*). Gross lesions were present predominantly in the middle, caudal and accessory lobes. Gross pathologically, lungs from 3D8 scFv pre-treatment group did not look different from the non-infected WT group. Marked pulmonary congestion and tan-yellow consolidation was observed in the PBS pre-treatment and the oseltamivir treatment groups (Fig. 5*A*). In the intergroup comparison, 3D8 scFv pre- and post-treatment groups had significantly low lung lesion scores compared with PBS and oseltamivir groups (*P* < 0.05). Between 3D8 scFv treatment groups, the gross lung lesion scores were significantly lower in the 3D8 scFv pre-treatment group than that of the 3D8 scFv post-treatment group (*P* < 0.05). The oseltamivir group had significantly lower gross lung lesion scores than PBS groups (*P* < 0.05) (Fig. 5*B*)

**Figure 5.**
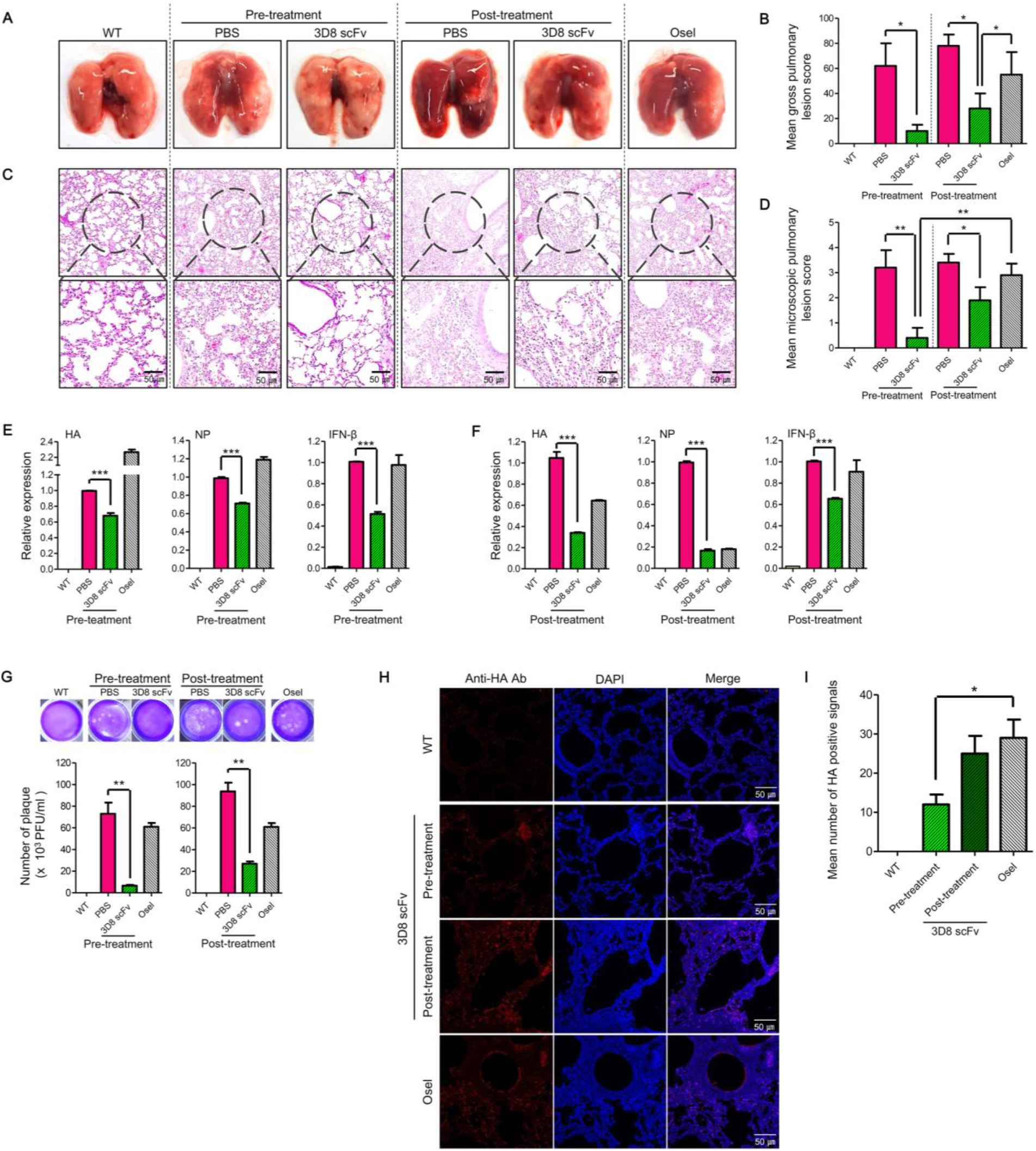
Histopathological analysis of lung tissues of H1N1/H275Y-infected mice after intra-nasally 3D8 scFv treatment. (*A* and *B*) Morphological comparison and gross pulmonary lesion score of mice lungs. Mice lungs were harvested at 5 dpi. (*C* and *D*) Histopathological comparison and Microscopic pulmonary lesion score of mice lung sections. Images of lung sections from virus-infected mice treated with 3D8 scFv. Lung tissues were stained with H&E. Scale bars represent 100 μm. (*E* and *F*) Viral RNA (HA, NP) and expression of the inflammatory factor, IFN-β were analyzed by qRT-PCR. The relative concentrations of RNA were calculated after normalization to GAPDH expression using delta Ct method. Data are shown as mean ± S.E.M of triplicate samples. (*G*) Comparison of virus titer in lung by a plaque assay. (*H*) Reduction in viral protein (HA) in infected lung detected using immunohistochemistry. Nuclei were detected using DAPI (Blue), and viral HA was visualized with TRITC (Red). Scale bars represent 50 μm. (*I*) Mean number of HA positive signal of mice lung. Error bars indicate SE. Asterisks indicate significant difference determined by *t*-test (*P < 0.05, **P < 0.01, ***P < 0.001).

Microscopical lesions were characterized by thickened alveolar septa with increased numbers of interstitial macrophages and lymphocytes in PBS and oseltamivir treatment groups. The lungs of 3D8 scFv pre-treatment and WT groups were normal (Fig. 5*C*). In the intergroup comparison, the 3D8 scFv pre-treatment group had significantly lower microscopic lesion scores than the other groups (*P* < 0.01) and the 3D8 scFv post-treatment group had significantly lower microscopic lesion scores than PBS and oseltamivir groups (*P* < 0.05). Microscopic lung lesion scores were not significantly different among PBS and oseltamivir groups (Fig. 5*D*).

Consistently, the expression of viral RNAs and IFN-β, an important pro-inflammatory mediator, at post viral infection was diminished in the lungs of the 3D8 scFv pre-treatment group compared with other groups, as demonstrated by reduced expression of HA (68%), NP (71%) and IFN-β (51%) in the lungs of the 3D8 scFv pre-treatment group compared with the PBS pre-treatment group (Fig. 5*E*). Moreover, the virus titer in the lung was reduced to 10% in the 3D8 scFv pre-treatment group, compared with the PBS pre-treatment group (Fig. 5*G*). Consistently, the signal of viral HA was reduced in the alveolar epithelial cell lining (Fig. 5*H*), and the number of distinct positive signals were significantly lower in the 3D8 scFv pre-treatment group compared with the oseltamivir group (P < 0.05; Fig. 5*I*).

In the 3D8 scFv post-treatment group, the expression of viral RNA decreased to 35% (HA) and 18% (NP) (Fig. 5*F*). Moreover, the virus titer in the lung was reduced to 30% in the 3D8 scFv post-treatment group, compared to the PBS post-treatment group (Fig. 5*G*). These results suggest that 3D8 scFv benefits virus-infected hosts elicited prophylactically as well as therapeutically by reducing viral burden, and alleviating excessive inflammation and potentially further pathogenesis.

## Discussion

Oseltamivir, an unrivaled antiviral treatment against IAV, has known to be effective in the early stages of viral infection by blocking NA, which is involved in progeny virus release (19). However, the H275Y missense mutation on NA gene in H1N1/pdm09 strain allosterically changes its binding affinity to the oseltamivir, resulting in 1,500-fold less sensitive to oseltamivir (20–22). Antigenic shifts and/or drifts are typical evolving strategies often selected by IAV, which has been major obstacle to develop anti-viral therapeutics (23, 24). The demand for pan-antiviral therapeutics that can exert a common effect regardless of the genetic mutation of IAV is increasing than ever (25).

In this study, we demonstrated that 3D8 scFv has antiviral effect in both oseltamivir-sensitive (H1N1/PR8 and H3N2/X-31) and oseltamivir-resistant (H1N1/H275Y) IAV that harbors a mutation, through its intrinsic viral RNA hydrolyzing activity. We report that, on an average, the *in vitro* cell viability rate increased by 20%, while the viral RNA expression reduced by about 50% within 24 h compared to the positive control in both 3D8 scFv pre- or post-treatment settings examined (Fig. 2 and 3). Consistently, *in vivo* intranasal administration of 3D8 scFv following viral infection elicited benefits as demonstrated by improved survival rate and bodyweight loss (Fig. 4). Viral infection after 3D8 scFv pre-treatment led to an increase in the survival rate from 37.5% and 30% in the PBS pre-treatment and oseltamivir groups, respectively, to 90%. The body weight decreased by 18% but recovered on 7 dpi (Fig. 4 *C* and *D*). Comparative assessment of the clinical indications (lung morphology, lung tissue sections, viral RNA/Protein/inflammation factors and number of plaque) among treatment groups revealed that 3D8 scFv pre-treatment showed a strong antiviral effect compared to PBS pre-treatment or oseltamivir group with minimal cytotoxicity (Fig. 5). This is because, during 48 h, 3D8 scFv was localized in the epithelial lining of bronchiole/alveoli such that when the cells are subsequently infected with the virus, the pre-existing 3D8 scFv starts to mediate the antiviral effects at the early stages of the infection (Fig. 6 and *SI Appendix,* Fig. S2). Based on that, 3D8 scFv can potentially be used as a prophylactic drug for influenza.

**Figure 6.**
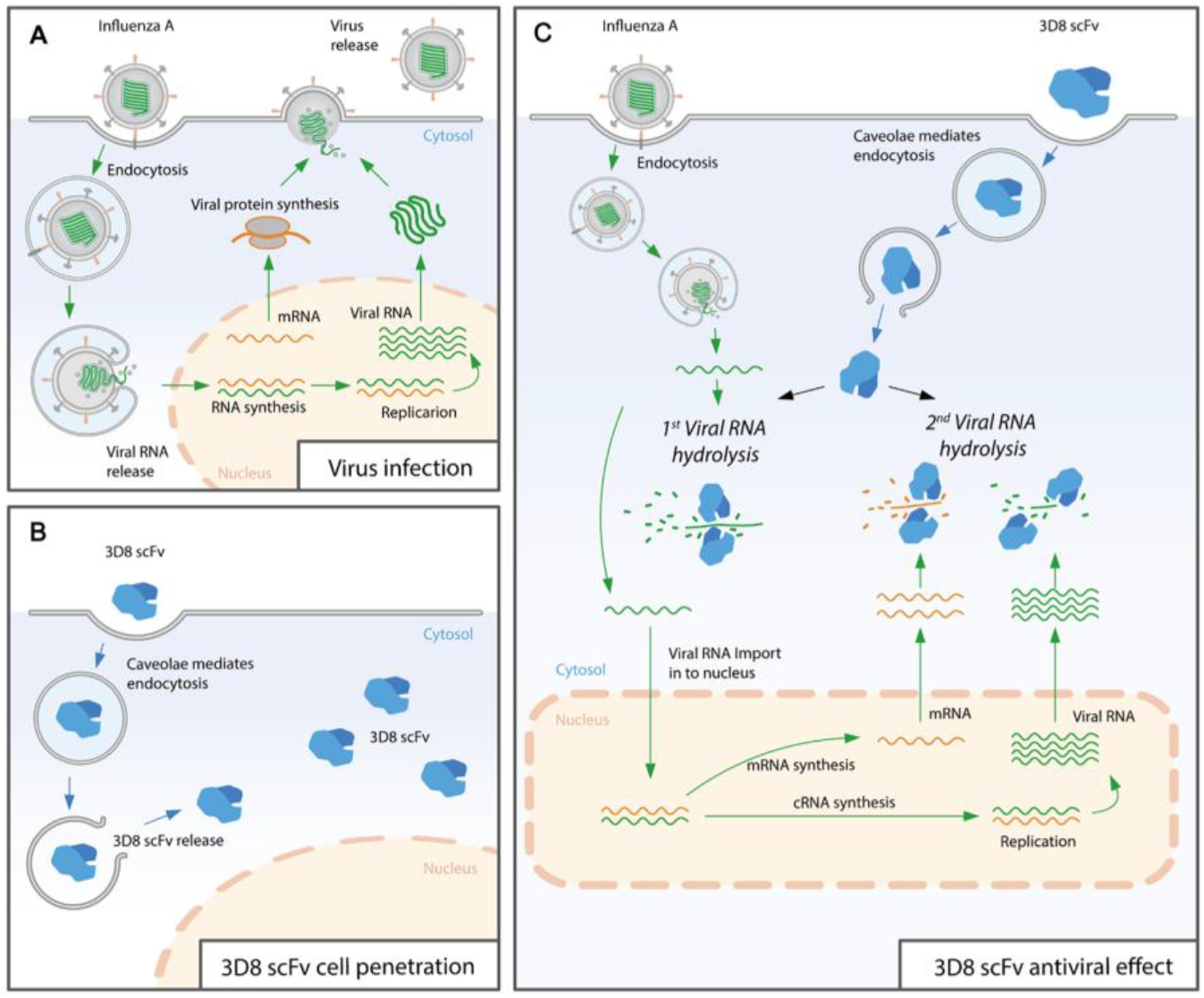
Proposed model of antiviral mechanism of action of 3D8 scFv. (*A*) Influenza virus replication pathway. (*B*) Cell penetration mechanism of 3D8 scFv. (*C*) Hydrolysis of influenza A virus RNA by 3D8 scFv in the cytosol.

When 3D8 scFv is administered intranasally, it is possible to passes through the respiratory mucosal surface and penetrates into the epithelial cells of the bronchiole/alveoli (*SI Appendix*, Fig. 2*D*). The mucosal surface is a selectively permeable barrier that covers the surface of internal organs and prevents the passage of toxins and bacteria (26, 27). The mucosal layer covers the respiratory (nose, trachea and lung) and digestive (intestine) epithelial cells. For the body to absorb a drug, it must pass through these layers (28). The above results imply that 3D8 scFv can potentially be used to manage infections of organs including respiratory tracts with a mucosal layer. For the nose-to-brain delivery of a drug through the nasal absorption route, the drug must pass through the mucosal layer of the nasal cavity first (29). Given that 3D8 scFv is capable of passing through the mucosal layer, it may be an important candidate for treating diseases resulting from infection of the central nervous system with the virus.

We also investigated the therapeutic effect of 3D8 scFv treatment after virus infection, to evaluate its potential as a drug that could be administered intranasally when the pathogenesis upon viral infection initiates and develops. Intranasal administration of PBS, which serve as control, after viral infection resulted in a survival rate of 0% at 8 dpi (Fig. 4*E*). However, the survival rate in the 3D8 scFv post-treatment group was 40%, and the body weight was restored after 10 dpi (Fig. 4*F*). Comparison of the clinical symptoms revealed that 3D8 scFv post-treatment showed antiviral effects compared to PBS post-treatment group (Fig. 5). Following viral infection, even when administered intranasally, 3D8 scFv exerts as an effective therapeutic drug.

Furthermore, 3D8 scFv may exhibit potent anti-viral effects against other RNA viruses with similar infection mechanisms to that of influenza virus, which potentially broaden the range of application to other virus types (Fig. 6). Due to 3D8 scFv is bio-production, there are many ways that can be applied genetically depending on the expression host. In our previous studies, we showed antiviral effects against H9N2 and newcastle disease using the 3D8 scFv transgenic chicken model (30, 31). Moreover, we also demonstrated the murine norovirus antiviral effect in 3D8 scFv-expression lactobacillus administration *in vivo* (32). In addition, we reconfirmed that 3D8 scFv expressed in recombinant lactobacillus did not induce any cytotoxic effect by NGS analysis (33). Based on the previous experiments in different mouse and chicken models against pseudoravis virus, influenza virus, norovirus and newcastle disease virus, 3D8 scFv can be a potentially broad spectrum antivirus protein. Although rarely observed in preclinical models, further systemic and careful investigation of toxicity and immune-related adverse events upon 3D8 scFv treatment should be performed to evaluate and predict the clinical efficacy of 3D8 scFv as anti-viral drug for patients.

In conclusion, 3D8 scFv can be applied as a potential prophylactic/therapeutic drug against the WT/mutated influenza virus because it directly affects viral RNA and blocks viral replication with non-sequence specific manner. The employment of 3D8 scFv will allow quicker response against new influenza outbreaks as well as newly emerging known or unknown viruses because processes such as the analysis of viral characteristics and identification of viral antigens can be avoided during the development of new therapeutic drugs for the novel influenza strain.

## Materials and Methods

### Cell lines and viruses

MDCK cells were maintained in Eagle’s minimal essential medium (EMEM) containing 5% fetal bovine serum (Gibco, Cergy Pontois, France), 100 U/ml penicillin and 100 μg/ml streptomycin (Hyclone, Logan, UT, USA). HeLa and Vero cells were grown in Dulbecco’s modified Eagle’s medium (DMEM), and A549 cells were cultured in Roswell park memorial institute 1640 medium (RPMI 1640 medium; Hyclone) containing the same supplements as the MDCK culture medium. The cell lines were purchased from the Korean Cell Line Bank and were maintained at 37℃ with 5% CO_2_. The influenza strain A/Puerto Rico/8/34 (H1N1/PR8) and A/X-31(H3N2/X-31) were kindly provided by Prof. Kweon (Sungkyunkwan University, Korea). Pandemic H1N1/H275Y NA-mutant virus (A/Korea/2785/2009pdm: NCCP 42017; Oseltamivir-resistant) were obtained from the National Culture Collection for Pathogens. The viruses were grown in the allantoic sacs of 9-day-old embryonated eggs at 37℃ for 3 days. The allantoic fluid was harvested and cleared using sucrose gradient centrifugation. Viral titer was determined using the plaque assay.

### Expression and purification of 3D8 scFv

3D8 scFv was expressed using pIg20-3D8 plasmid-transformed *Escherichia coli* (BL21[DE3]*pLysE*) strain in Luria-Bertani (LB) broth with 100 μg/ml ampicillin and 20 μg/ml chloramphenicol at 37℃, and induced for 18 h at 26℃ by adding 1 mM isopropyl 1-thiol-β-D-galactopyranoside (IPTG). The cell culture supernatant was obtained by centrifugation at 5,200 × g for 20 min at 4℃ and was pass through a 0.22 μm filter. 3D8 scFv was purified from the supernatant using an IgG Sepharose 6 fast flow (GE Healthcare, Malborough, MA, USA) affinity column. The column was washed with ten bed volumes of PBS (pH 7.4) and then with ten bed volumes of 5 mM ammonium acetate (pH 5.0). Next, 3D8 scFv was eluted with 0.1 M acetic acid (pH 3.4). The eluted fraction was neutralized with 0.1 volume of 1 M Tris-HCl (pH 9.0). The concentration of purified 3D8 scFv was determined based on the extinction coefficient at 280 nm. The protein was analyzed by electrophoresis on a 12% SDS polyacrylamide gel and detected by Coomassie blue staining.

### Preparation of NA and NP RNA transcripts and analysis of RNA hydrolyzing activity of 3D8 scFv

Total RNA was isolated from H1N1/PR8 and viral cDNA was synthesized. The sequences corresponding to NA and NP were amplified using PCR (Table S1) and cloned into pGEM-T Easy, a vector that harbors T7 and SP6 promoter (Promega, Madison, WI, USA). NA and NP RNA were synthesized using a HiScribe T7 *in vitro* transcription kit (New England BioLabs, Ipswich, MA, USA) and incubated with 3D8 scFv purified protein (0.5 μg) for 1 h in TBS containing 0.1 mM MgCl_2_ at 37℃. The reaction was terminated by adding 10× loading buffer (Takara, Otsu, Shinga, Japan) and analyzed by electrophoresis on a 1% agarose gel and stained with ethidium bromide.

### Analysis of cell penetration in response to 3D8 scFv using immunocytochemistry (ICC)

A549, HeLa, Vero, and MDCK cells (2 × 10^4^) were cultured in 8-well chamber slides (Nunc Lab-Tek). Cells were treated with 3 μM 3D8 scFv and incubated at 37℃ with 5% CO_2_ for 24 h. Next, the slides were washed with PBS twice and fixed for 15 min in chilled methanol at 25℃. The samples were permeabilized for 5 min using Intracellular Staining Perm wash buffer (BioLegend, San Diego, CA, USA). After blocking with 1% BSA in PBST for 1 h, the samples were incubated with monoclonal mouse anti-3D8 scFv antibody (1:1000 dilution; AbClon, Seoul, Korea) at 4℃ for 24 h. Next, the samples were incubated with goat anti-mouse IgG Alexa Fluor 488 secondary antibody (1:1000 dilution; Abcam, Cambridge, MA, USA) for 1 h at 25℃ and the slides were mounted in antifade Mounting Medium with DAPI (Vectashield, Burlingame, CA, USA). The samples were then visualized using a Zeiss LSM 700 confocal microscope.

### Analysis of cytotoxicity of 3D8 scFv

MDCK cells (2 × 10^4^) were cultured in 96-well plates. Cells were treated with 3D8 scFv (1–10 μM) and incubated at 37℃ with 5% CO_2_ for 48 h. Then, 10 μl of MTT solution was added and the cells were incubated at 37℃ with 5% CO_2_ for 4 h. The formazan crystals formed were dissolved with DMSO and absorbance of each well was read at 595 nm using an ELISA microplate reader (TECAN, Mannedorf, Switzerland).

### Analysis of the antiviral activity (cell viability test) of 3D8 scFv

MDCK cells (2 × 10^4^) were cultured in 96-well plates and treated as follows: (ⅰ) 3D8 scFv pre-treatment: cells were treated with 3D8 scFv (3, 5, and 7 μM) at 37℃ with 5% CO_2_ for 24 h and then the cells were independently infected with the three influenza viral strains (MOI = 0.1) in MEM-free media for 1 h at 37℃. Then the infection media was removed and the cells were cultured in MEM-free media (1% BSA) containing tosyl phenylalanyl chloromethyl ketone (TPCK)-treated trypsin (1 μg/ml) at 37℃ with 5% CO_2_ for 48 h. (ⅱ) 3D8 scFv post-treatment: MDCK cells were independently infected with the three influenza virus strains (MOI = 0.1) in MEM free media for 1 h at 37℃ followed by removal of the infection media. The medium was changed to MEM-free media (1% BSA) containing 3D8 scFv (3, 5, and 7 μM) and TPCK-treated trypsin (1 μg/ml) followed by incubation at 37℃ with 5% CO_2_ for 48 h (Fig. 2w *A* and *E*). Then 10 μl of MTT solution was added to each well and the cells were incubated at 37℃ with 5% CO_2_ for 4 h. The formazan crystals formed were dissolved with DMSO and the absorbance at 595 nm was measured using an ELISA microplate reader.

### Measurement of the expression of viral RNA using qRT-PCR

MDCK cells (4 × 10^5^) were cultured in 6-well plates and treated as follows: (ⅰ) 3D8 scFv pre-treatment: cells were treated with 3 μM of 3D8 scFv at 37℃ with 5% CO_2_ for 24 h and then infected independently with the three influenza viral strains (MOI = 0.1) in MEM-free media for 1 h at 37℃ followed by the removal of the infection media. The medium was changed to MEM-free media (1% BSA) containing TPCK treated trypsin (1 μg/ml) followed by incubation at 37℃ with 5% CO_2_ for 24 h. (ⅱ) 3D8 scFv post-treatment: MDCK cells were infected with the three influenza virus strains (MOI = 0.1) in MEM-free media for 1 h at 37℃ followed by the removal of the infection media. The medium was changed to MEM-free media (1% BSA) with 3D8 scFv (3 μM) containing TPCK-treated trypsin (1 μg/ml) followed by incubation at 37℃ with 5% CO_2_ for 24 h (Fig. 3*A*). Total RNA was isolated using TRI reagent (MRC, Montgomery, OH, USA) according to the manufacturer’s instructions. RNA concentration was determined using a spectrophotometer. cDNA was synthesized using CellScript All-in-One 5× First Strand cDNA Synthesis Master Mix (Cell safe, Yongin, Korea) according to the manufacturer’s protocol. Quantitative real-time PCR was performed using SYBR Premix Ex Taq and Rotor-Gene Q system. Data were analyzed using Rotor-Gene Q series software version 2.3.1 (Qiagen, Chadstone, Victoria, Australia). The genes (HA and NP) were amplified using the indicated primers (Table S2). GAPDH was amplified as an internal control.

### Measurement of viral HA expression using ICC

MDCK cells (2 × 10^4^) were cultured in 8-well chamber slides. The procedures for 3D8 scFv treatment and virus infection (H1N1/PR8 and H1N1/H275Y) were same as described in the previous section (Fig. 3*A*). Next, the slides were washed with PBS twice and fixed for 15 min using chilled methanol at 25℃. Samples were permeabilized for 5 min with Intracellular Staining Perm wash buffer. After blocking with PBST containing 1% BSA for 1 h, the cells were incubated with monoclonal mouse anti-3D8 scFv antibody or polyclonal rabbit anti-HA antibody (1:1000 dilution; Invitrogen) for 24 h at 4℃. Then the cells were incubated with goat anti-mouse IgG Alexa Fluor 488 secondary antibody or goat anti-rabbit IgG TRITC secondary antibody (1:1000 dilution; Abcam), respectively, for 1 h at 25℃. The slides were mounted using anti-fade Mounting Medium with DAPI and the cells were visualized using a Zeiss LSM 700 confocal microscope.

### Animals and antiviral activity test in vivo

Six-week-old male specific pathogen-free (SPF) BALB/c mice (DBL; weighing 18–20 g) were housed under standard laboratory conditions. All animal procedures were approved by the Institutional Animal Care and Use Committee of Sungkyunkwan university (Permit number: SKKUIACUC2019-03-07-3). Mice were pre-/post-treated intranasally with 50 μg of 3D8 scFv for 4 days. Oseltamivir phosphate (Sigma-Aldrich, ST. Louis, MO, USA) was administered orally for 5 days post infection. The mice were infected intranasally with 50 μl lethal dose 50% of H1N1/H275Y virus (5 × 10^4^ PFU; Fig. 4 *A* and *B*). After challenge with H1N1/H275Y, the mice were monitored daily for clinical signs (survival and weight loss) until day 11 post-infection. Lung samples were collected for histopathological examination on 5 dpi.

### Measurement of viral RNA and interferon levels

Total RNA was extracted from lysed mouse lung tissue (5 dpi) samples using TRI reagent according to the manufacturer’s instructions. The RNA concentration was determined using a spectrophotometer. cDNA was synthesized using CellScript All-in-One 5× First Strand cDNA Synthesis Master Mix according to the manufacturer’s protocol. The genes (HA, NP, and IFN-β) were amplified with the indicated primers (Table S2) using qRT-PCR as described previously. GAPDH was amplified as an internal control.

### Comparison of virus titer in lung

Mouse lung tissues were homogenized in 1 ml of MEM-free media using TissueRuptor (Qiagen, Chadstone, Victoria, Australia). The homogenate was obtained by centrifugation at 6,000 × g for 5 min at 4℃. Confluent MDCK cells were cultured in 12-well plates and then infected with the tissue supernatant (1:1000 dilution) in MEM-free media for 1 h at 37℃ followed by the removal of the infection media. The media was replaced with 1× Dulbecco’s modified eagle medium containing TPCK treated trypsin (1 μg/ml) and 1% agarose. The cells were further incubated at 37℃ with 5% CO_2_ for 3 days. The cell monolayers were fixed with 4% formaldehyde and stained with 0.5% crystal violet for visualization.

### Immunohistochemistry (IHC)

Tissue sections were deparaffinized, rehydrated, and antigen retrieval was performed using citrate buffer at 95℃ for 20 min. Slide were washed in TBST (TBS, 0.025% Triton X-100) and blocked using a solution of 10% fetal bovine serum + 1% BSA in TBST for 2 h. The tissue sections were incubated with polyclonal rabbit anti-HA antibody (1:750 dilution) overnight at 4℃. The tissue slides were then incubated for 1 h at 25℃ with goat anti-rabbit IgG TRITC secondary antibody (1:1000 dilution), respectively. The tissue sections were finally mounted in anti-fade Mounting Medium with DAPI and the cells were visualized using an LSM 700 Zeiss confocal microscope.

### Morphometric analysis

Lung tissues were collected from each group of mice. For morphometric analysis of the gross pulmonary lesion score, the method described elsewhere by Harbur *et al*.(1995) was slightly modified (34). Briefly, each lung lobe was assigned a number that reflected the approximate percentage of the volume of the entire lung represented by that lobe. The total points for all of the lobes formed an estimate of the percentage of the entire lung with grossly visible pneumonia.

For morphometric analysis of the microscopical pulmonary lesion score, lung tissues were fixed in 10% neutral buffered formalin. After fixation, the tissues were dehydrated through a graded series of alcohol solutions and xylene and embedded in paraffin wax. Sections (5 μm) were prepared from each tissue and followed by haematoxylin and eosin (HE) staining. Prepared tissue slides were examined in blinded fashion and scored for the estimated severity of the interstitial pneumonia as: 0, no lesions; 1, mild interstitial pneumonia; 2, moderate multifocal interstitial pneumonia; 3, moderate diffuse interstitial pneumonia; and 4, severe interstitial pneumonia.

For morphometric analysis of IHC, the sections were analyzed with the NIH Image J 1.43m Program (https://imagej.nih.gov/ij) to obtain quantitative data. Ten fields were selected randomly and the number of positive cells per unit area (0.25 mm_2_) was determined. The mean values were then calculated.

### Statistical analysis

Continuous data were analyzed with a one-way analysis of variance (ANOVA). If a one-way ANOVA was significant (*P* < 0.05), pairwise Tukey’s adjustment was performed as a posthoc. Discrete data were analyzed by Chi-square and Fisher’s exact test. If necessary, Pearson’s correlation coefficient and/or Spearman’s rank correlation coefficient was used to assess the relationship among data. A value of *P* < 0.05 was considered significant. All statistical analyses were performed using the GraphPad Prism software (GraphPad Software, San Diego, CA, USA).

## Acknowledgments

We thank Drs. Song, Minkyung and Jeon, Youngjun for editing the manuscript. This work was supported by a grant from Novelgen Company (Project NO. S-2018-1158-000).

