## Supplement Figure 1, Supplement Figure 2, Supplement Table 1, Supplement Table 2 for "Antiviral effect of a nucleic acid hydrolyzing scFv against oseltamivir resistant influenza A virus"

**This PDF file includes:**

Supplementary Materials and Methods

Figures S1 and S2

Tables S1 and S2

### Materials and Methods

#### Cell lines and viruses

MDCK cells were maintained in Eagle's minimal essential medium (EMEM) containing 5% fetal bovine serum (Gibco, Cergy Pontois, France), 100 U/ml penicillin and 100 μg/ml streptomycin (Hyclone, Logan, UT, USA). The cell lines were purchased from the Korean Cell Line Bank. The cells were maintained at 37℃ in 5% $CO_2$ atmosphere. The influenza strain A/Puerto Rico/8/34 (H1N1/PR8) and A/X-31(H3N2/X-31) were kindly provided by Prof. Kweon (Sungkyunkwan University, Korea). Pandemic H1N1/H275Y NA-mutant virus (A/Korea/2785/2009pdm: NCCP 42017; Oseltamivir resistance) was obtained from the National Culture Collection for Pathogens. The viruses were grown in the allantoic sacs of 9-day-old embryonated eggs at 37℃ for 3 days. The allantoic fluid was harvested and cleared using sucrose gradient centrifugation. Viral titer was determined using the plaque assay.

#### Oseltamivir antiviral activity test (cell viability)

MDCK cells ($2 \times 10^4$) growing in 96-well plates were infected with H1N1/PR8, H3N2/X-31, or H1N1/H275Y (MOI = 0.1) in MEM free media for 1 h at 37℃, and then the infection media was removed and replaced with MEM free media (1% BSA) containing oseltamivir phosphate (Sigma-Aldrich, ST. Louis, MO, USA) at various concentrations (1/2 serially diluted starting from 500 μM) containing tosyl phenylalanyl chloromethyl ketone (TPCK)-treated trypsin (1 μg/ml). The cells were incubated at 37℃ in 5% $CO_2$ for 48 h and then treated with 10 μl of MTT solution at 37℃ in 5% $CO_2$ for 4 h. The formazan crystals were dissolved in DMSO and the absorbance at 595 nm was measured using an ELISA microplate reader (TECAN, Mannedorf, Switzerland).

#### Animals and safety test in vivo

Six-week-old male specific pathogen-free (SPF) BALB/c mice (DBL; weighing 18–20 g) were housed under standard laboratory conditions. All animal procedures were approved by the

Institutional Animal Care and Use Committee of Sungkyunkwan university (Permit number: SKKUIACUC2019-03-07-3). Mice were treated intranasally with 50 μg of 3D8 scFv for 4 days (Supplementary Fig. 2a). Lung samples were collected daily for histopathological examination until day 5 after administration.

**Histopathological staining**

Lung tissues were collected from each group of mice and were fixed in 10% neutral buffered formalin, embedded in paraffin wax and sectioned at 5-µm. The tissue sections were mounted on slides, stained with H&E for examination, and scanned using slide scanner Axioscan Z1.

**Immunohistochemistry (IHC)**

Tissue sections were deparaffinized, rehydrated, and antigen retrieval was performed using citrate buffer at 95℃ for 20 min. Slide were washed in TBST (TBS, 0.025% Triton X-100) and blocked using a solution of 10% fetal bovine serum + 1% BSA in TBST for 2 h. The tissue sections were incubated with polyclonal rabbit anti-3D8 scFv antibody (AbFrontier, Seoul, Korea) overnight at 4℃. The tissue samples were then incubated for 1 h at 25℃ with goat anti-rabbit IgG Alexa Fluor 488 secondary antibody (1:1000 dilution). The tissue sections were finally mounted in anti-fade Mounting Medium with DAPI and the cells were visualized using an LSM 700 Zeiss confocal microscope.

**Measurement of 3D8 scFv proteins retention time.**

Total protein was extracted from homogenized mouse lung tissue samples using Pro-Prep solution (iNtRON bio, Seongnam, Korea) according to the manufacturer's instructions. Protein concentration was determined using the Bradford assay. 3D8 scFv was analyzed through electrophoresis on a 12% SDS gel, and transferred onto a PVDF membrane (GE Healthcare, Malborough, MA, USA) using a semi-dry method. The membrane was incubated with polyclonal rabbit anti-3D8 antibody or polyclonal rabbit anti β-actin antibody (1:1000 dilution) at 4℃ for 24 h.

Then the membrane was incubated with a goat anti-rabbit IgG-HRP secondary antibody (1:5000 dilution) at 25℃ for 1 h. The size of bands was measured using ImageJ software.

**Statistical analyses**

All statistical analyses were performed using the GraphPad Prism software (GraphPad Software). One-way ANOVA was used for statistical analyses. All data are presented as the mean ± standard error (SE).

**Supplementary Figures and Tables**

A

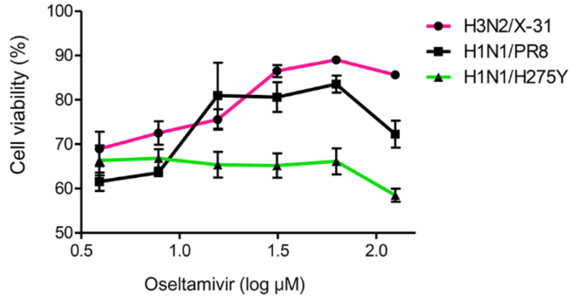


**Supplementary Figure 1. Antiviral activity of oseltamivir against the three influenza virus stains.** (*A*) Viability of virus-infected MDCK cells following oseltamivir treatment. MDCK cells were infected with viruses (MOI = 0.1) for 1 h followed by the removal of the viral inoculum. The medium was replaced with new media containing various concentrations of oseltamivir (serially diluted [1/2] in PBS with a starting concentration of 500 μM). After incubation for 48 h at 37℃, cell viability was measured using the MTT assay. Data are shown as mean ± S.E.M of triplicate samples.

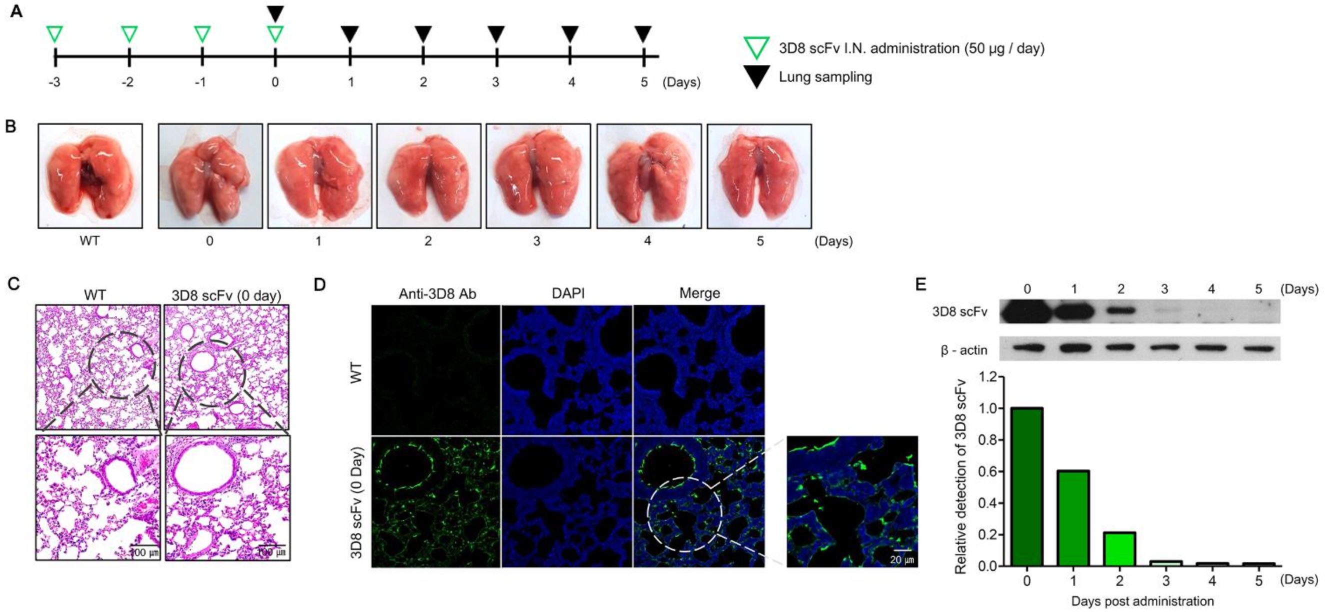


**Supplementary Figure 2. Evaluation of the pharmacological properties of 3D8 scFv *in vivo*.** (*A*) Schematic diagram of 3D8 scFv treatment. 3D8 scFv (50 μg/day) was administered intranasally to mice for 4 days. (*B*) Morphological comparison of mice lungs. Lungs were harvested every day for 5 days. (*C*) Histopathological comparison of H&E-stained lung sections from mice treated with 3D8 scFv. Scale bars represent 100 μm. (*D*) 3D8 scFv penetration in lung tissue was detected using immunohistochemistry. Nuclei and 3D8 scFv were detected using DAPI (blue) and Alexa flour 488 (Green). Scale bars represent 50 μm. (*E*) 3D8 scFv retention time in mouse lungs was analyzed using western blotting.

**Supplementary Table 1. Specific primers used in PCR amplification of viral (A/PuertoRico/8/1934 [H1N1]) transcripts**

| Gene name | Forward (5' – 3') | Reverse (5' – 3') | Accession No. |
|---|---|---|---|
| Neuraminidase (NA) | CCA TTG GAT CAA TCT GTC TGG | TGT CAA TGG TGA ATG GCA ACT C | NC_002018.1 |
| Nucleoprotein (NP) | ACT GAT GGA GAA CGC CAG AAT | ATG TCA AAG GAA GGC ACG ATC | NC_002019.1 |

**Supplementary Table 2. Specific primers used in qRT-PCR**

| Gene name | Forward (5' – 3') | Reverse (5' - 3') | Accession No. |
|---|---|---|---|
| GAPDH (MDCK Cell Line) | AAC ATC ATC CCT GCT TCC ACT | GGC AGG TCA GAT CCA CAA C | NM_001003142.2 |
| GAPDH (Mouse) | GGC ATT GCT CTC AAT GAC AA | TGT GAG GGA GAT GCT CAG TG | NC_000072.7 |
| Hemagglutinin (A/PuertoRico/8/1934 (H1N1)) | AGT GCC CAA AAT ACG TCAG G | CAG TCC ATC CCC CTT CAA TA | NC_002017.1 |
| Hemagglutinin (A/X-31 (H3N2)) | GAA AAT GGT TGG GAG GGA ATG | GAT GGC TGC TTG AGT GCT TTT | DQ874876.1 |
| Hemagglutinin (A/California/07/2009 (H1N1)) | GGA ACG TGT TAC CCA GGA G | GCT GCC GTT ACA CCT TTG TT | NC_026433.1 |
| Nucleoprotein (A/PuertoRico/8/1934 (H1N1)) | ACC AAT CAA CAG AGG GCA TCT | TGA TTT CGG TCC TCA TGT CAG | NC_002019.1 |
| Nucleoprotein (A/X-31 (H3N2)) | AAA ATC ATG GCG TCT CAA GGC | CGG TGC ACA TTT GGA TGT AGA | AB036779.1 |
| Nucleoprotein (A/California/07/2009 (H1N1)) | CCA GAT CAG TGT GCA GCC TA | GGC TTT GCA CTT TCC ATC ATT C | NC_026436.1 |
| Interferon beta (Mouse) | TTA CAC TGC CTT TGC CAT CCA A | TCC CAC GTC AAT CTT TCC TCT T | NC_000070.6 |
